# Influence of bacterial swimming and hydrodynamics on infection by phages

**DOI:** 10.1101/2024.01.15.575727

**Authors:** Christoph Lohrmann, Christian Holm, Sujit S. Datta

**Affiliations:** Institute for Computational Physics, University of Stuttgart, 70569 Stuttgart, Germany; Department of Chemical and Biological Engineering, Princeton University, Princeton, New Jersey 08544, USA

## Abstract

Bacteriophages (“phages”) are viruses that infect bacteria. Since they do not actively self-propel, phages rely on thermal diffusion to find target cells—but can also be advected by fluid flows, such as those generated by motile bacteria themselves in bulk fluids. How does the flow field generated by a swimming bacterium influence how it encounters and is infected by phages? Here, we address this question using coupled molecular dynamics and lattice Boltzmann simulations of flagellated bacteria swimming through a bulk fluid containing uniformly-dispersed phages. We find that while swimming increases the rate at which both the cell body and flagellar propeller are infected by phages, hydrodynamic interactions strongly *suppress* this increase at the cell body, but conversely *enhance* this increase at the flagellar bundle. Our results highlight the pivotal influence of hydrodynamics on the interactions between bacteria and phages, as well as other diffusible species in microbial environments.

## I. INTRODUCTION

More than 10^31^ bacteriophages (“phages” for short) are estimated to exist on Earth, more than every other organism on the planet combined [1, 2]. Indeed, on average, phages—viruses that infect bacteria—outnumber bacterial cells by a factor of ten [3]. Thus, their interactions with phages regulate how bacteria function in natural environments, with critical implications for agriculture, ecology, and medicine [4–8]. Extensive research has focused on documenting various biological and chemical factors that influence these interactions; nevertheless, the basic physical processes underlying how bacteria encounter and thereby become infected by phages in the first place remain poorly understood.

For example: while phages do not actively self-propel, many bacteria do using, e.g., flagella-driven swimming through fluids. We therefore focus on this mode of motility here. How does swimming influence the rate at which bacteria encounter and become infected by phages, if at all? Despite its apparent simplicity, this question still does not have a clear answer [9]—even for the illustrative, idealized case of a cell swimming at a constant velocity *v*_swim_ in an unbounded, uniform, Newtonian fluid at low Reynolds number. One might expect that the rate at which the cell is infected by phages simply increases proportionately with its swimming speed as it explores more space [10]; however, this expectation does not consider the complex flow field generated by the bacterium around itself as it swims, which can advect surrounding phages in a non-trivial manner [11]. The importance of these hydrodynamics is highlighted by examining the Péclet number comparing the characteristic rates of phage transport by fluid advection and thermal diffusion, Pe_phage_ ≡ *v*_swim_*r*_body_*/D*^P^, which exceeds unity for typical values of the bacterial swimming speed *v*_swim_ ∼ 10 − 100 μm s^*−*1^, size *r*_body_ ∼ 1 μm, and phage diffusivity *D*^P^ ∼ 1 −10 μm^2^ s^*−*1^. Hence, the rate at which a bacterium encounters and becomes infected by phages is likely highly sensitive to the nature of the flow field it generates by swimming.

In a classic study [12], Berg and Purcell examined the influence of these hydrodynamics by treating the swimming bacterium as an externally-driven sphere. Building on prior calculations [13, 14], the authors concluded that because the cell pushes fluid around it as it moves, “motility cannot significantly increase the cell’s acquisition of material”, and the phage infection rate only increases sublinearly with swimming speed. However, while instructive, this analysis neglects two crucial features of flagellated bacteria—which are not uniform spheres, but are typically comprised of an elongated cell body driven by an adjoined elongated flagellar propeller. First, it is now well known that many such bacteria (including *Escherichia coli, Bacillus subtilis*, and *Salmonella enterica*) are force-free “pushers”: each cell pushes on surrounding fluid with an equal and opposite force to the one generated by its flagella as it swims [11, 15, 16]. As a result, the fluid boundary conditions are distinct for the cell body and the flagella, and phage infection rates may thus differ considerably between the two. Second, the flow field away from a swimming bacterium has a dipolar shape with a fluid velocity magnitude *v* that decays with distance *r* away from the cell as ∼ 1*/r*^2^, unlike the longer-ranged *v*(*r*) ∼ 1*/r* decay characteristic of a driven sphere. How these two features influence the manner in which swimming bacteria encounter and become infected by phages is as yet unknown.

Here, we address this gap in knowledge using simulations of a “pusher”-type bacterium swimming through a fluid containing uniformly-dispersed phages. We use particle-resolved molecular dynamics to explicitly treat the cell body, flagellar propeller (hereafter referred to as the flagellum for convenience), and individual phages, all of which are coupled to and interact via an underlying fluid which we simulate using a lattice Boltzmann algorithm. We find that while swimming increases the rate at which the cell body is infected by phages, the dipolar flow thereby generated advects phages away from the forward-facing “head” of the cell body, thereby suppressing this increase by as much as twofold—as suggested by Berg and Purcell. However, this dipolar flow field also pumps phage-containing fluid to the flagellum. As a result, the phage infection rate at the flagellum is appreciable and increases nearly linearly with swimming speed, in stark contrast to the findings of Berg and Purcell. Altogether, our work suggests that while the fluid flow generated by swimming helps to protect the bacterial cell body from phages, it *promotes* phage infection at the flagellum—an effect that, to the best of our knowledge, has been overlooked in previous work. This effect may be exploited by flagellotropic phages, i.e., phages that initiate infection by attaching to the host flagellum, which are increasingly being recognized as key constituents of natural microbial communities and potentially useful therapeutics against pathogenic bacteria [17, 18]. More broadly, our findings highlight the pivotal influence of hydrodynamics on the interactions between bacteria and phages, as well as other diffusible species in microbial environments.

## II. METHODS

### A. Molecular dynamics model of phages and bacterium

We use a particle-resolved, coarse-grained molecular dynamics algorithm to represent the phages and bacterium. Particles of each are coupled to and interact with each other via an underlying lattice Boltzmann fluid through friction forces, as detailed below. While here we focus on the case of a Newtonian fluid, which is relevant to many aquatic microbial habitats, studying the case of a more complex non-Newtonian fluid [10, 19–30] will be a useful extension of our work.

#### Phages

For simplicity, we treat phages as uniform spherical particles. Their dynamics in three spatial dimensions are described by the Langevin equations

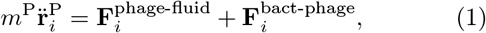

where

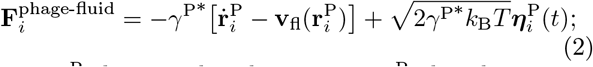

here, *m*^P^ denotes the phage mass, **r**^P^ the phage position, *i* the particle index, the overdot a derivative with respect to time, 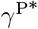 the phage-fluid friction coefficient, **v**_fl_ the fluid velocity, *k*_B_ Boltzmann’s constant, *T* the temperature, ***η***^P^ a noise term, and **F**^bact-phage^ the phage-bacterium interaction force. The noise term has mean ⟨ ***η***^P^ ⟩= **0** and correlations 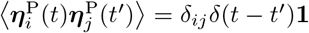, where ⟨·⟩ denotes an ensemble average and **1** the identity matrix in three dimensions. Our focus here is on phage-bacteria encounters, and we therefore do not consider any phage-phage interactions.

We treat hydrodynamics at a continuum level by solving the Navier-Stokes equations using a multi-relaxation time lattice Boltzmann algorithm with Guo forcing [31]. Thermal fluctuations of the fluid are included following the approach of Ref. [32]. To obtain the fluid velocity **v**_fl_ at the particle positions 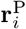, we interpolate between the nearest lattice nodes according to Ref. [33]. The coupling force **F**^phage-fluid^ reflects the microscopic interactions between the fluid and the phage particles. These interactions obey Newton’s third law, so the coupling force is also applied to the fluid, but with opposite sign. We thus extrapolate − **F**^phage-fluid^ to the nearest lattice node as detailed in Ref. [33].

#### Bacterium

Our model for the bacterium builds on previous studies of microswimmers [34–37], extended to also explicitly consider the cell and flagellum geometry. Specifically, we represent the bacterium by a rigid collection of three different types of particles denoted by *τ* to model the rod-like shape of flagellated bacteria like *E. coli, B. subtilis*, and *S. enterica*. The first type (large green spheres in Fig. 1) consists of spheres that make up the cell body. They experience point-friction with the fluid and interact with phage particles via an almost-hard-sphere potential that defines the cell body volume, as detailed further below. The second type (smaller green spheres in Fig. 1) consists of points placed on the surface of particles of the first type that only experience friction with the fluid, thereby approximating a no-slip boundary condition on the surface of the cell body. The third type (medium blue spheres in Fig. 1) consists of spheres that make up the flagellum. Similar to the first type, they interact with phages via an almost-hard-sphere potential. However, unlike those comprising the cell body, these particles do not experience friction with the fluid. Instead, they exert a force on the fluid that is equal in magnitude but opposite in sign to the swim force **F**^swim^ that propels the cell body forward, ensuring that the overall swimming bacterium is net force-free. This model represents a coarse-grained picture of a swimming bacterium that does not take into account the helical geometry of a flagellum or the intricate inner structure of a flagellar bundle. Rather, it seeks to capture four essential features of the bacterial flagellar bundle: it is permeable to fluid, exerts a force on the fluid, occupies a non-zero volume, and crucially, that phages can attach to it.

**FIG. 1.**
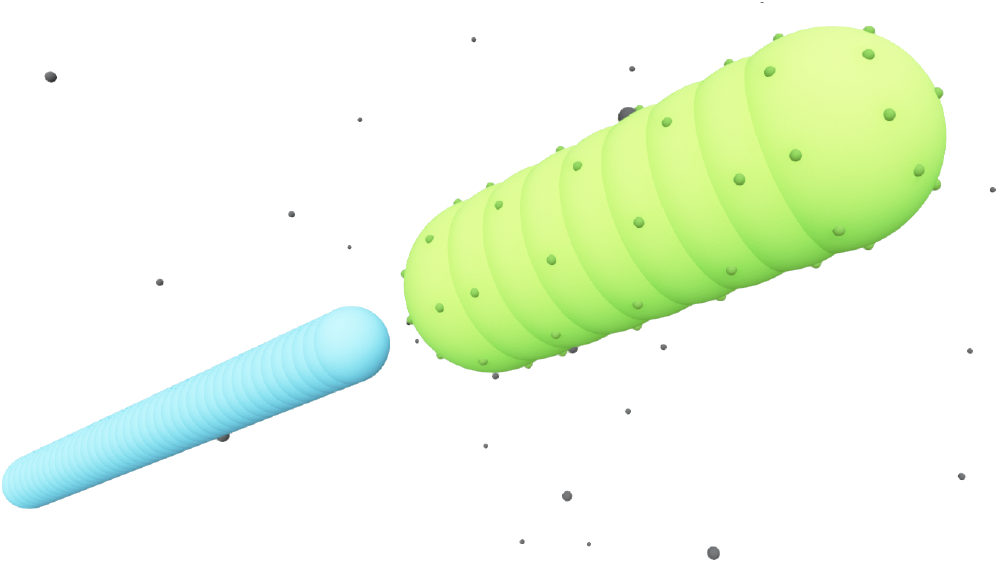
Rendering of our rigid-body, coarse-grained molecular dynamics model of a swimming bacterium. Large green and medium blue spheres comprise the cell body and flagellum, respectively. Small, darker green spheres mark the locations of particle-fluid coupling points. Small gray spheres represent phages dispersed in the surrounding fluid.

More specifically, the cell body comprises *N* ^body^ particles of type one arranged in a line along **ê**, evenly spaced over the length *l*_body_, along with *N* ^coupl^ additional coupling points of type two on the surface of the cell body. The dynamics of the central particle of the cell body in three spatial dimensions are described by the equations of motion

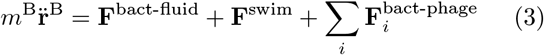

for the center of mass position and

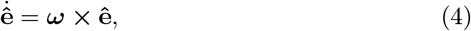

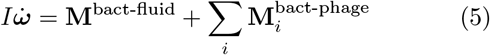

for the orientation; here, *m*^B^ denotes the mass of the cell, **r**^B^ its position, **F**^swim^ ≡ *F* ^swim^ **ê** the swim force of magnitude *F* ^swim^ and orientation **ê, *ω*** the angular velocity, *I* the moment of inertia tensor, and the summation runs over all phage particles *i*. All other particles add their forces and torques to the central particle, themselves following its trajectory in their predefined relative position and orientation to keep the shape of the rigid bacterium. The resulting coupling force is given by

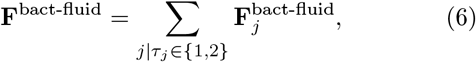

where the summation runs over all particles *j* with type *τ*_*j*_ associated with the cell body. The individual coupling forces are calculated analogous to Eq. (2):

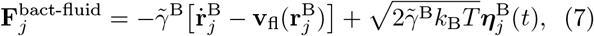

where 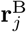 denotes the position of the *j*th particle of the cell body, 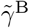 the bare friction coefficient, and 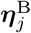 a noise term with the same properties as ***η***^P^. As for the phages, all coupling forces 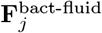 are also applied to the fluid in the opposite direction to ensure net momentum conservation. The coupling forces also lead to the torque

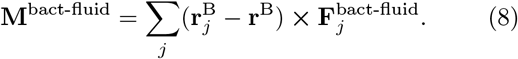

The flagellum is modelled by *N* ^flagellum^ particles of type three arranged in a line along **ê** and evenly spaced over a length *l*_flag_, with the central particle of the flagellum at a distance *l*_dipole_ behind the central particle of the cell body. Flagellum particles do not experience friction or noise as the other particles in the system. Instead, each of them applies a force

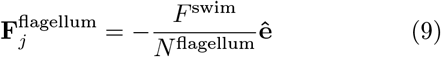

to the fluid, such that the forward force on the bacterium is balanced by a backward force on the fluid and the total force resulting from bacterial swimming is zero.

Cell body particles of type one and flagellum particles interact with phages via the short-ranged, purely-repulsive, almost-hard-sphere Weeks-Chandler-Anderson potential [38]

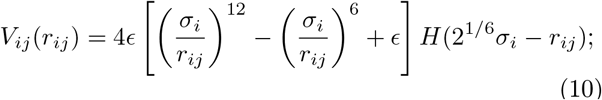

here, 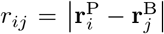 is the distance between bacterium particle *j* and phage *i, ϵ* the interaction strength, and *H*(·) the Heaviside step-function. We choose *ϵ* = 10*k*_B_*T* to limit overlap between particles. The scale parameter *σ*_*i*_ depends on the types of particles involved. For cell body particles of type one, it is given by 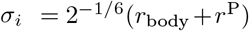, while for flagellum particles, it is given by 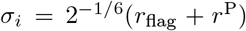, where *r*_body_, *r*_flag_, *and r*^P^ are the radii of the cell body, the flagellum, and the phages, respectively. Forces and torques are then given by the gradient of this potential:

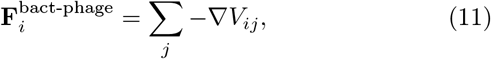

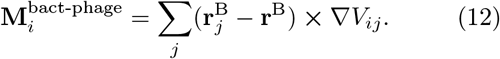

In our simulations, we solve Eqs. (1)–(12) numerically using the development version of the ESPResSo simulation package [39] with the waLBerla [40, 41] library for lattice Boltzmann hydrodynamics. Exact software versions as well as our custom code are freely available; details are provided in [42].

#### Simulations without hydrodynamics

As a further test of the influence of hydrodynamics, we also perform simulations that do not include the hydrodynamic field. For the phage particles, we set **v**_fl_(**r**^P^) = **0**. For the bacterium, the equations further simplify, because we do not introduce the additional coupling particles of type two. Instead, we have 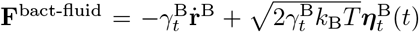 and 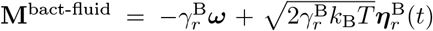 where 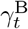 and 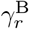 are the translational and rotational friction coefficients, respectively.

### B. Model of phage infection

We consider a phage as being in contact with the bacterium (either the cell body or flagellum) if the surface-to-surface distance is smaller than a prescribed encounter distance *d*_enc_. Once it contacts the bacterium, the phage attaches and infects the cell at a rate *k*^att^, which is a lumped parameter that describes the complex process of binding to specific receptors [43, 44]. As we show below, the overall qualitative insights that result from our simulations are insensitive to the specific choice of *k*^att^. In our model, this attachment step completes the infection; because our focus is on the purely physical processes underlying phage encounter and infection, we do not consider the subsequent biological steps needed for the phage to actually insert its genetic material into the cell.

In our time-discretised numerical implementation, we check for phages in contact with the bacterium after every successive time interval Δ*t*. For such contacting phages, we calculate the attachment probability *p*^att^ = Δ*tk*^att^. Based on this probability, using a random number generator, we choose whether the phage remains free or it successfully attaches and infects the cell. If infection happens, we register the time and relative position of the phage encounter on the bacterium surface. We then move the phage to a random position outside the encounter region defined by *d*_enc_ to prevent double-counting; this procedure mimics the case of a bacterium swimming in an infinite fluid reservoir with constant phage number density. We then determine the overall infection rate 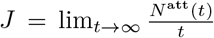 from a linear fit to the computed number of phages that have infected the cell at time *t, N* ^att^(*t*) [45]. For the case of *k*^att^ →0 (*p*^att^→ 0), the number density of phages in the encounter region is asymptotically independent of *k*^att^. Hence, *J* scales linearly with *k*^att^, and we only need to determine the number density of phages in the encounter region. Therefore, for simulations of *k*^att^→ 0, we compute *J* by tracking the trajectories of phages in the contact region to obtain their number density.

### C. Continuum modelling

To compare the results of our particle-based simulations to those of Berg and Purcell [12], we follow their approach and solve the continuum advection-diffusion equation

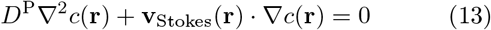

for the phage concentration *c*. As in their work, we model the bacterium as a perfectly-absorbing sphere of radius 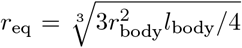 in an infinite reservoir of phages, with boundary conditions *c*(|**r**| = *r*_eq_) = 0 and *c*(|**r**| → ∞ = *c* _∞_ The Stokes flow field is given by

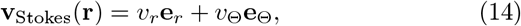

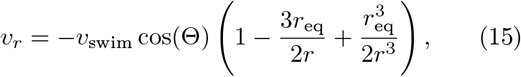

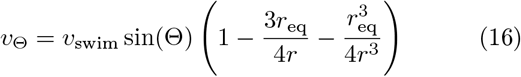

in the comoving frame of reference; here, *r* and Θ denote the radius and polar angle in spherical coordinates, and **e**_*r*_, **e**_Θ_ the corresponding basis vectors. The infection rate is then given by the flux of phages into the absorbing sphere surface *Ω*_*s*_; we compute it directly from the solution of the concentration field via the relation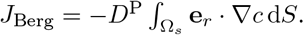

In practice, we solve Eqs. (13)–(16) using a finite element algorithm [46] with the boundary condition *c*( |**r**| = *R*) = *c*_0_ with *R* = 50 μm ≫*r*_eq_. The finite size effects from restricting the simulation domain to this radius are negligible, and the infection rate for *v*_swim_ = 0 obtained with finite *R* deviates from the analytical solution for *R* → ∞ only by ≈1 %.

### D. Choice of numerical parameters

For the phages, we choose *r*^P^ = 50 nm, which is comparable to the size of many commonly-studied phages, including T4 [47] and the flagellotropic phages *χ* and PBS1 [18]. We calculate the phage friction coefficient from Stokes’ law as *γ*^P^ = 6*πμ*_water_*r*^P^, where *μ*_water_ = 1× 10^*−*3^ Pa· s is the dynamic shear viscosity of water; the numerical friction coefficient then follows from a lattice correction given by 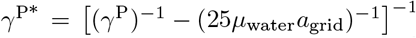, where *a*_grid_ is the lattice Boltzmann grid constant [33]. The phage diffusion coefficient is given by the Einstein-Smoluchowski relation *D*^P^ = *k*_B_*T/γ*^P^ ≈ 4.4 μm^2^ s^*−*1^ at our choice of *T* = 300 K, which is in good agreement with experimental measurements [10, 48]. We calculate the phage mass as *m*^P^ = *ρ*^P^4*/*3*π*(*r*^P^)^3^, where *ρ*^P^ is the mass density of the phage. If this density is taken to be that of water, *ρ*_water_ = 1000 kg m^*−*3^, the time scale for momentum relaxation 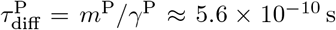 s is much smaller than any time scale of interest in our system. The dynamics are therefore overdamped and the exact value of *m*^P^ does not influence our results. Thus, we instead choose *ρ*^P^ = 10^5^*ρ*_water_, which leaves the dynamics still overdamped but allows us to use a larger numerical integration time step *δt* = 3.33 ×10^*−*5^ s.

To mimic the commonly-studied flagellated bacterial species *E. coli* [49] and *S. enterica* [50], we choose *l*_body_ = 3 μm and *r*_body_ = 0.5 μm, with *N* ^body^ = 9 and *N* ^coupl^ = 62 to obtain a sufficiently well-resolved cell surface. The bacterial mass is approximated by 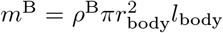 as detailed above for the phages, the diffusive time scale for bacteria is very small and the dynamics are over-damped, so we choose *ρ*^B^ = 5 × 10^3^*ρ*_water_. For simulations without hydrodynamics, we use a spheroidal approximation, with the translational and rotational friction coefficients 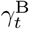 and 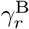 calculated following Refs. [51] and [52], respectively. The force needed to propel the bacterium with the desired velocity is then given by 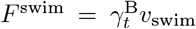. For simulations with hydrodynamics, we set 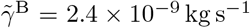, which represents the largest value that we can use without impeding numerical stability to most closely approximate a no-slip boundary condition. The relation between *F* ^swim^ and *v*_swim_ is non-trivial because of the complex flow field and the many coupling points; however, empirically, we find that using *F* ^swim^ = *γ*^eff^*v*_swim_ with an effective friction coefficient 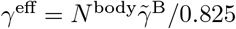 yields good agreement between the desired and actual swimming velocity, as detailed further in [45]. The flagellum is *l*_flag_ = 5 μm long with a radius of *r*_flag_ = 0.25 μm at a distance *l*_dipole_ = 5 μm, and we take *N* ^flagellum^ = 41. For comparison to other kinds of microswimmers, we also perform all our simulations with a puller-type swimmer with *l*_dipole_ = − 5 μm. Those results are presented in [45].

The lattice Boltzmann fluid has a dynamic viscosity *μ* = *μ*_water_ and is discretised on a grid with lattice spacing *a*_grid_ = *r*_body_. Since all velocities are interpolated between lattice nodes, this resolution is fine enough to resolve the flow near the bacterium surface. To keep simulations stable, we choose a fluid mass density of *ρ*_fluid_ = 1200*ρ*_water_. This choice does not change the flow behaviour with respect to water since the Reynolds number Re ≡ *ρ*_fluid_*v*_swim_*r*_body_ */μ*≈ 0.017 for a typical swimming velocity *v*_swim_ = 25 μm s^*−*1^ is much smaller than unity, even for this increased density.

Our simulations consider one bacterium and *N* ^P^ = 1000 phage particles in a fully periodic, cubic simulation domain with box length *L* = 16 μm. The volume fraction of phages *ϕ* = *N* ^P^4*π*(*r*^P^)^3^*/*3*L*^3^ ≈ 10^*−*4^ is very low, so we expect any hydrodynamic interactions between them to be negligible. Moreover, in the supplementary material [45] we present evidence that there are no appreciable finite-size effects associated with the size of the simulation domain. We set the encounter distance *d*_enc_ = *r*^P^ = 50 nm; the time scale for phage diffusion across this distance is 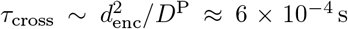, so we choose the Δ*t* = 1 ×10^*−*4^ s to adequately register phage encounters.

Each simulation is repeated *N* ^ensemble^ times with different seeds, i.e., different initial conditions and noise realisations, to obtain the standard error of the mean of the quantities reported below. We choose *N* ^ensemble^ ≥5, the exact numbers for each reported result can be found in the data set [42] that accompanies this work.

## III. RESULTS

Before considering interactions with phages, we first verify that our simulation approach recapitulates the farfield dipolar flow field characteristic of swimming flagellated bacteria [11, 15]. This is indeed the case, as shown in Fig. 2, which shows the fluid flow field in the laboratory frame of reference. Importantly, while the bacterium moves in the positive *z*-direction, our model for the flagellum-fluid interaction leads to fluid motion in the negative *z*-direction inside the flagellum, with swimming being force-free overall.

**FIG. 2.**
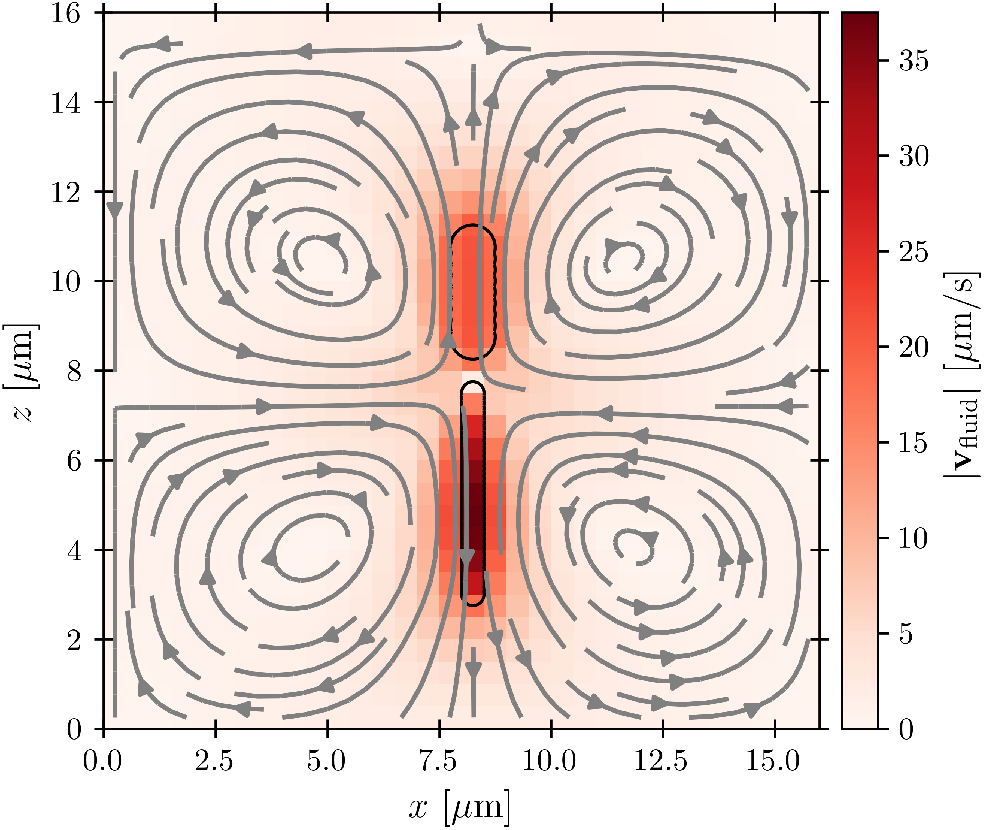
Cross-section through the dipolar flow field generated by a swimming bacterium, in the stationary lab frame of reference. This representative example is for the case of *v*_swim_ = 20 μm s^*−*1^ in the absence of thermal noise in the lab frame. Color scale and arrows show the magnitude and orientation of the local fluid velocity, respectively. The upper and lower black outlines show the cell body and flagellum, respectively; the cell is swimming in the +*z* direction.

Having established that our simulations reproduce this characteristic dipolar flow field, we next investigate what its implications are for infection by phages. As described below, we use our simulations to examine phage interactions first with the cell body, then with the flagellum.

### A. Phage infection of the cell body

To characterize how bacterial swimming and the associated hydrodynamic interactions (HI) influence how phages infect the cell body, we compute the infection rate *J* over a broad range of representative swimming speeds *v*_swim_, for a broad range of *k*^att^, with or without HI included. Our results are presented in Fig. 3 for the limiting cases of *k*^att^→ 0 and *k*^att^ → ∞for clarity; the intermediate values of *k*^att^ monotonically interpolate between these cases, as shown in the supplementary material [45].

**FIG. 3.**
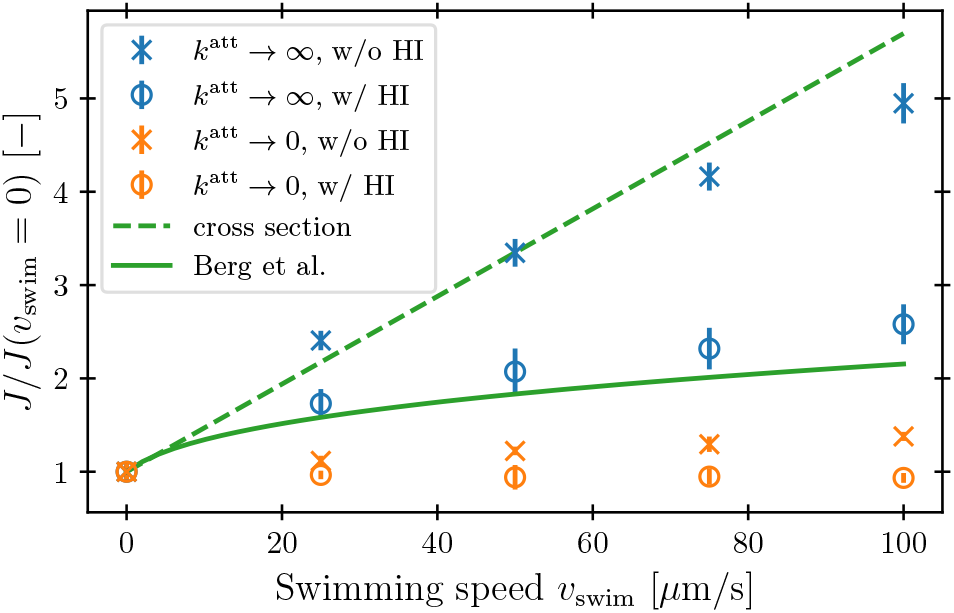
Hydrodynamic interactions (HI) suppress the increase in phage infection rate on the cell body with swimming speed. The infection rate *J* integrated over the cell body is computed from our simulations and normalized by its value when the bacterium is not swimming. Blue and orange points show the cases of rapidly- or slowly-attaching phages, respectively. Error bars represent the standard error of the mean over statistically independent simulations. Dashed green line corresponds to a model of rapid phage infection through diffusion and uptake by the cell body cross section, neglecting HI. Solid green line shows the prediction of Berg and Purcell from a more simplified continuum model of a driven spherical bacterium.

We first consider the case of rapidly-attaching phages (*k*^att^ → ∞). In the absence of HI, *J* increases approximately linearly with *v*_swim_ (blue ×, Fig. 3), as suggested previously [10]. Indeed, without HI, we expect that 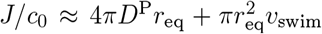; here, the first term on the right hand side describes the flux of a diffusive species into a spherical, stationary, perfect absorber [53] and is given by the solution of Eq. 13 with *v*_swim_ = 0, and the second term takes the motion of the absorber into account by calculating the rate at which its cross section explores the volume filled with a constant number density of phages that are absorbed upon contact. As shown by the dashed green line in Fig. 3, this prediction agrees well with our simulation results.

Incorporating HI strongly suppresses infection by phages, however. While *J* still increases monotonically with *v*_swim_, the magnitude of this increase is considerably lessened by hydrodynamics (blue ◦, Fig. 3)—indeed, by as much as twofold at the largest swimming speed tested. Why is this the case?

Close inspection of the flow field around the cell body provides a clue: as shown in Fig. 2, the no-slip boundary condition on the cell body surface causes a region of fluid in its immediate vicinity to be dragged along with it as it swims. We expect that this region also advects surrounding phages along with the cell, pushing them away from its forward-facing “head”; consequently, these phages must rely primarily on passive thermal diffusion to cross this region and successfully infect the cell, independent of the bacterial swimming speed.

This effect is more clearly visualized in the reference frame that moves along with the bacterium. The relative flow field given by 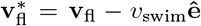, where the asterisk denotes quantities measured in this comoving frame, is shown in Fig. 4. We quantify the relative importance of phage advection and diffusion using the local Péclet number 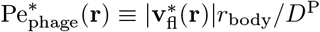 defined in this co-moving frame; the iso-line of 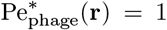 is shown by the dashed line. As expected, within the region bordered by the dashed line, the fluid is nearly at relative rest and 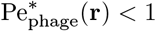—confirming that any phages contained therein must rely primarily on thermal diffusion to infect the cell body. Any enhancement in the phage infection rate arising from bacterial swimming is therefore suppressed by this “protective shield” of fluid that surrounds the cell body.

**FIG. 4.**
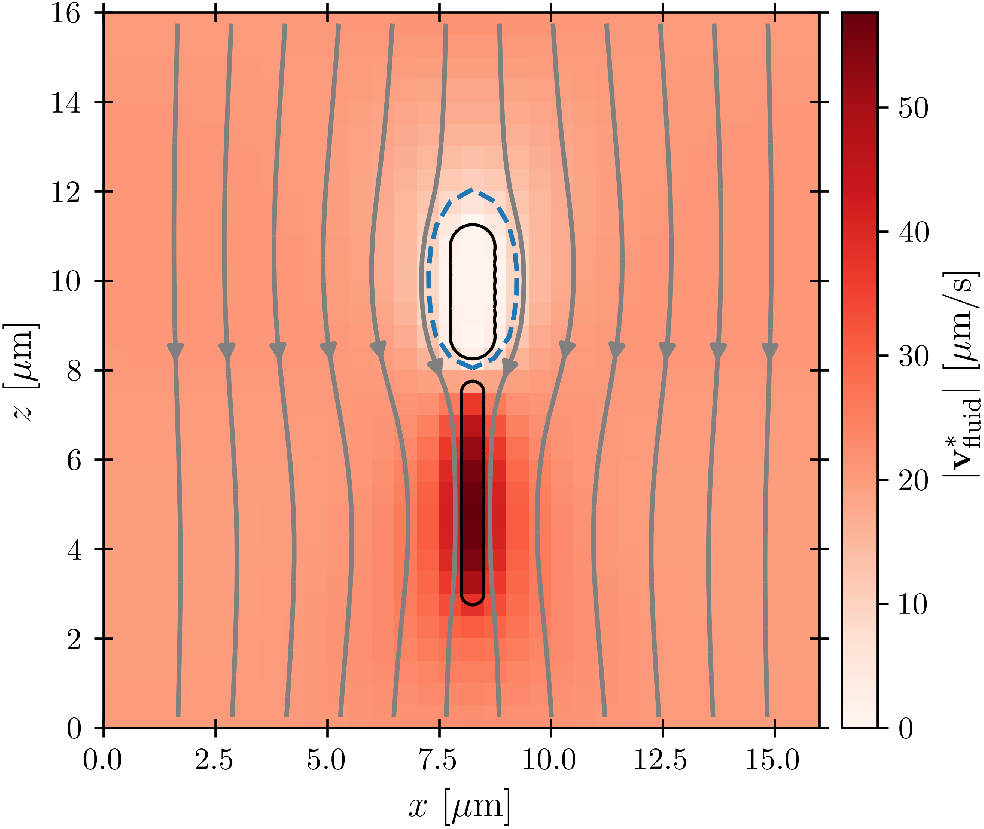
Cross-section through the dipolar flow field generated by a swimming bacterium, in the cell’s moving frame of reference. The image shows the same flow field as in Fig. 2, but in the comoving frame. The cell is swimming in the +*z* direction. The blue dashed line marks 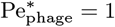.

This hydrodynamic effect manifests at smaller phage attachment rates (*k*^att^→ 0), as well. When HI are not included, the swimming bacterium is not protected by the “shield” of quiescent fluid discussed above. Instead, as it swims, the cell collects and accumulates phages at its head, and *J* again increases approximately linearly with *v*_swim_, as shown by the orange × in Fig. 3. In this case, the swimming bacterium does not appreciably remove contacting phages from the fluid, whereas in the case of *k*^att^ → ∞, phages are locally depleted from the vicinity of the cell upon contact and infection. Therefore, the dependence of *J* on *v*_swim_ is more modest for poorly-attaching phages; indeed, when HI are included, there is no measurable dependence of *J* on *v*_swim_ (orange ◦, Fig. 3).

The protective influence of the hydrodynamic “shield” is also apparent in the spatial distribution of phage infection over the cell body, shown for the example of *v*_swim_ = 100 μm s^*−*1^ and *k*^att^ →0 in Fig. 5. As expected, without HI (left panel), infections occur preferentially at the head of the cell. However, HI push phages away from the head of the cell, causing infections to be distributed more uniformly around the cell body (right panel). Fig. 6 quantifies this difference for all the simulation conditions tested using the average *z*-position of phage infection relative to the cell body center of mass, i.e., the first moment of the infection probability density. Again, as shown by the difference between × and ◦ symbols, HI reduce the asymmetry of phage infection for both *k*^att^ → 0 and *k*^att^ → ∞ over all swimming speeds tested.

**FIG. 5.**
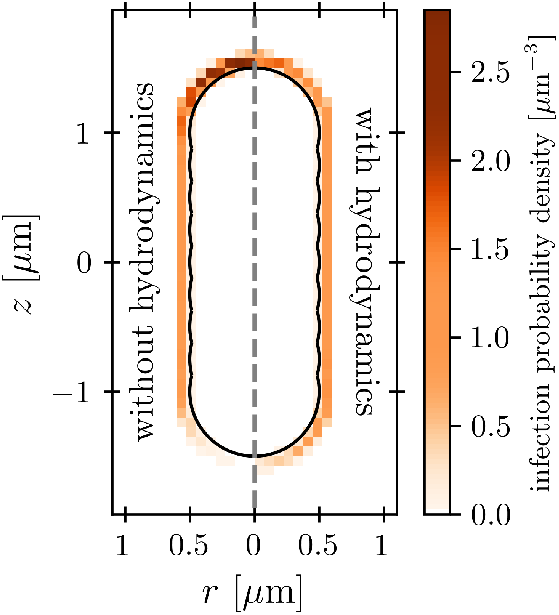
Hydrodynamic interactions cause phages to be advected away from the head of the cell. Color scale shows the probability density of phage infection over the surface of the cell body, for the example of *v*_swim_ = 100 μm s^*−*1^ and *k*^att^ *→* 0. The cell is swimming in the +*z* direction. With-out hydrodynamics, phages accumulate at its head, whereas when hydrodynamics are incorporated, the flow field established by swimming makes the distribution of infection sites more uniform along the cell body.

**FIG. 6.**
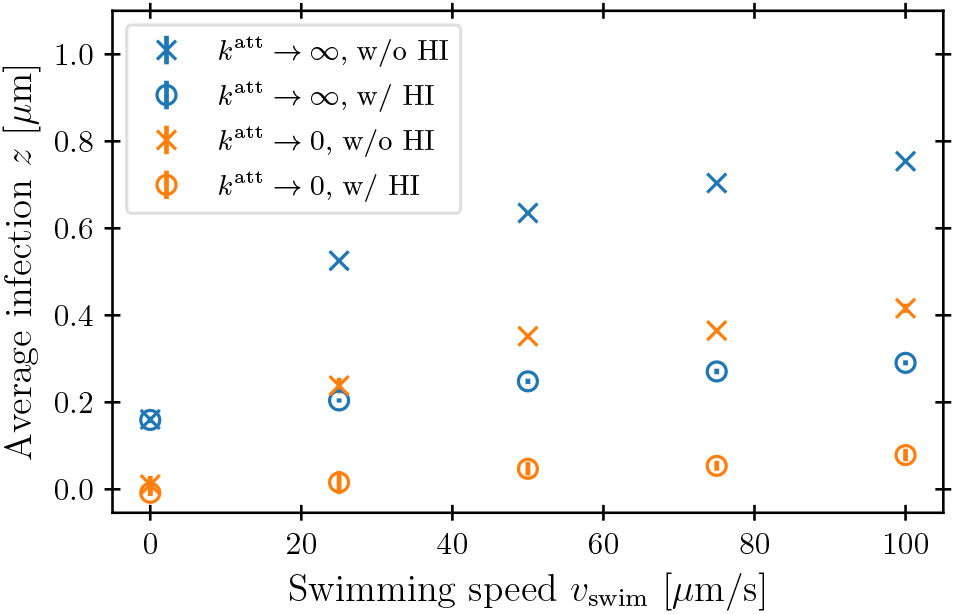
Hydrodynamic interactions cause phages to be advected away from the head of the cell. Symbols show the average position of phage infection positions along the bacterial symmetry axis relative to the cell body center. An average of *z >* 0 means the infection probability density is shifted towards the head of the cell. Blue and orange points show the cases of rapidly- or slowly-attaching phages, respectively. Error bars represent the standard error of the mean over statistically independent simulations, and are sometimes smaller than the symbol size.

In their classic paper [12], Berg and Purcell did not consider the elongated shape of a swimming bacterium, the distinction between the cell body and flagellum, or the dipolar flow field established by swimming. Nevertheless, using a more simplified continuum model of an externally-forced sphere, they intuited the protective hydrodynamic effect uncovered by our simulations, noting that “The molecules [or in our case, phages] in front of the cell are carried out of its way along with the fluid it must push aside to move. The cell carries with it a layer of liquid that is practically stationary in its frame of reference. Every molecule [or phage] that reaches the surface of the cell must cross this layer by diffusion.” Remarkably, despite the simplifications made in their work, Berg and Purcell’s prediction for the infection rate (solid green curve, Fig. 3) shows excellent agreement with the results of our more detailed simulations (blue ◦). One may not expect this agreement *a priori* given the marked difference between the actual dipolar flow field established by the swimming cell and the approximation used by Berg and Purcell. However, our results show that the essential qualitative feature of the low 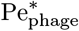 region are independent of the shape and propulsion mechanism of the bacterium, justifying their simplifying assumptions *a posteriori*—but only for the case of the cell body. As we show in the next section, we find dramatically different behavior for the bacterial flagellum.

### B. Phage infection of the flagellum

While the cell body drags a protective region of fluid with it, the flagellum does not. Instead, because it is permeable to and exerts a force on the fluid, 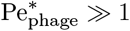 especially in the immediate vicinity of the flagellum, as seen by the dark red region of Fig. 4. As a result, we expect that hydrodynamic interactions are not protective as for the cell body, but instead, *promote* infection of the flagellum. Our simulations confirm this expectation, as shown in Fig. 7.

**FIG. 7.**
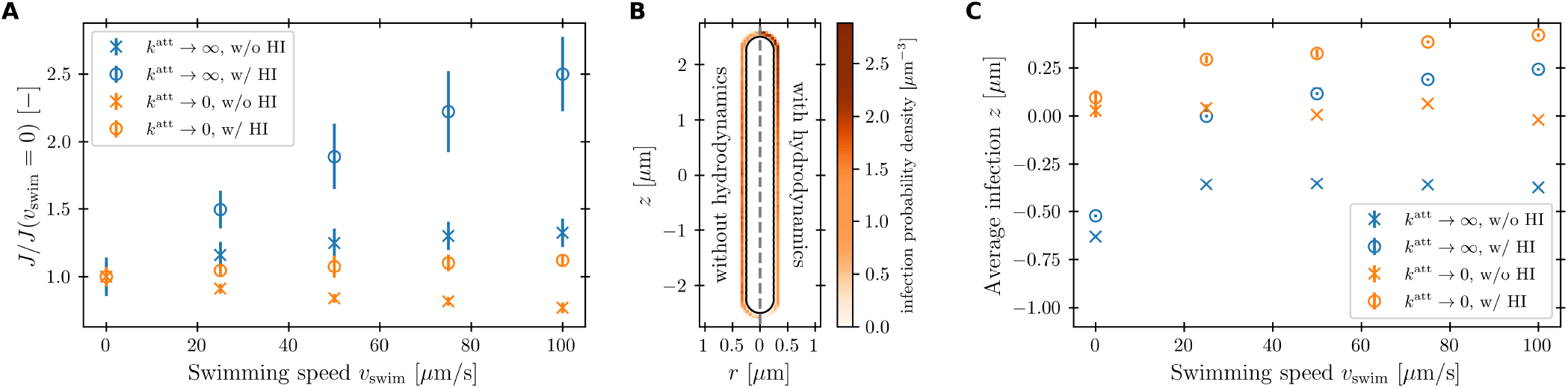
Hydrodynamic interactions promote phage infection of the flagellum. (A) The infection rate *J* integrated over the flagellum is computed from our simulations for varying *v*_swim_ and normalized by its value when the bacterium is not swimming. (B) Color scale shows the probability density of phage infection over the surface of the flagellum, for the example of *v*_swim_ = 100 μm s^*−*1^ and *k*^att^ *→* 0. The cell is swimming in the +*z* direction. When hydrodynamics are incorporated, fluid pumping by the flagellum along the *−z* direction drives phage infection near the forward end of the flagellum. (C) Symbols show the average position of phage infection positions along the bacterial symmetry axis relative to the flagellum center. An average of *z >* 0 means the infection probability density is shifted towards the forward end of the flagellum. For *k*^att^ *→ ∞*, the attachment is shifted into the negative direction at *v*_swim_ = 0 because the cell body depletes the region in front of the flagellum of phages. Error bars in A and C represent the standard error of the mean over statistically independent simulations. Blue and orange points show the cases of rapidly- or slowly-attaching phages, respectively.

Unlike the case of the cell body, HI greatly *increase* the flagellar infection rate, as seen by comparing the ◦ and × points in Fig. 7A—indeed, by nearly twofold at the largest swimming speed tested. In the absence of HI, *J* only increases marginally with *v*_swim_ for *k*^att^ → ∞. It even decreases with *v*_swim_ for *k*^att^ → 0, due to the cell body pushing phages radially outward upon contact. However, when HI are taken into account, the flagellum pumps in phage-laden fluid from the sides of the cell body and moves it through the space occupied by the flagellar bundle. Therefore, the volume of fluid coming in contact with the flagellum, and thus phage infection, increases with *v*_swim_. Moreover, because the fluid is pumped along the −**ê** direction, we expect that phage infection is more likely to occur at the forward end of the flagellum with increasing *v*_swim_. Inspecting the spatial distribution of phage infection along the flagellum confirms this expectation, as shown for the example of *v*_swim_ = 100 μm s^*−*1^ and *k*^att^→ 0 in Fig. 7B, and quantified for all the simulation conditions tested in Fig. 7C.

## IV. CONCLUSIONS

Using coarse-grained molecular dynamics simulations of a swimming bacterium that explicitly treat its cell body and flagellum separately, with hydrodynamic interactions incorporated via coupling to a lattice Boltzmann fluid, our work has shed new light on the influence of swimming on infection by phages. We find that while swimming increases the rate at which both the cell body and flagellar propeller are infected by phages, hydrodynamic interactions strongly *suppress* this increase at the cell body, but conversely *enhance* this increase at the flagellar bundle. This difference in infection arises from the characteristic dipolar flow field generated by a swimming bacterium, which advects phages away from the cell body, but pumps phage-laden fluid into the flagellum. Hence, while our results corroborate the findings of Berg and Purcell for the cell body, our work provides a counterpoint to their conclusion that “in a uniform medium motility cannot significantly increase the cell’s acquisition of material [in our case, phages].” Experimentally testing these predictions—e.g., by combining direct visualization of phage infection [54] with optical trapping of swimming cells [55]—will be an important direction for future work.

Altogether, our findings highlight the pivotal influence of hydrodynamics on the interactions between bacteria and phages, as well as other diffusible species—e.g., nutrients, toxins, or signalling molecules—in microbial environments. They also provide a new perspective on the biophysical tradeoffs associated with bacterial swimming. While swimming can be beneficial by enabling bacteria to escape from harmful environments, find new resources, and colonize new terrain, it can also be costly—not only because of the additional energy it requires of the cell, but also because it is often thought to increase the probability of encountering surrounding phages. Our results demonstrate that this latter cost is not appreciable for the cell body, due to the protective “shield” of fluid established by hydrodynamics, but is appreciable for the flagellar bundle, which pumps surrounding phages in. We conjecture that this may in part be why many phages have, over billions of years of evolution, developed ways to exploit bacterial swimming for their benefit by targeting the flagellum [17, 18]. Swimming bacteria may, in turn, have evolved localised defence countermeasures against phage infection such as through modification of specific surface receptors or production of outer membrane vesicles as decoys [56]. Investigating how these biological and chemical processes, combined with the hydrodynamic effects illuminated by our work, influence phage-bacteria interactions will be a useful avenue for future research.

## DATA AND CODE AVAILABILITY

The data that support the findings of this study as well as the source code are available in [42].

## AUTHOR CONTRIBUTIONS

C.L. performed all simulations and theoretical calculations; C.L., C.H., and S.S.D. designed the simulations, analyzed the data, discussed the results and implications, and wrote the manuscript; S.S.D. designed and supervised the overall project.

## ACKNOWLEDGEMENTS

We thank the Visiting Student Research Collaborator (VSRC) program of the Graduate School at Princeton for enabling C.L. to visit the Datta Lab and conduct this research, as well as funding support by the Deutsche Forschungsgemeinschaft (DFG, German Research Foundation) under Project Number 327154368-SFB 1313 and under Germany’s Excellence Strategy EXC 2075 - 390740016. The simulations were performed on computational resources managed and supported by Princeton Research Computing, a consortium of groups including the Princeton Institute for Computational Science and Engineering (PICSciE) and the Office of Information Technology’s High Performance Computing Center and Visualization Laboratory at Princeton University. It is a pleasure to thank Ido Golding and Ned Wingreen for useful discussions, and Chris Browne for helpful comments on the manuscript.

## Supplementary information

### I. BACTERIUM AND PHAGE MODEL CALIBRATION

Figure S1 shows that using our choice of 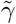 and determining the corresponding effective mobility leads to the desired swim speed of the bacterium. In fig. S2 we show that applying the grid correction to *γ*^*P*^ leads to the correct diffusion coefficient of the phages.

**FIG. S1.**
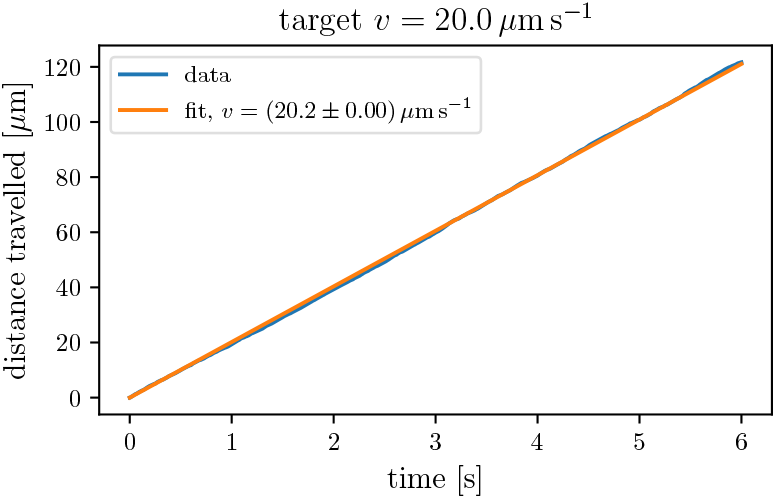
Bacterium distance travelled and swim speed fit.

**FIG. S2.**
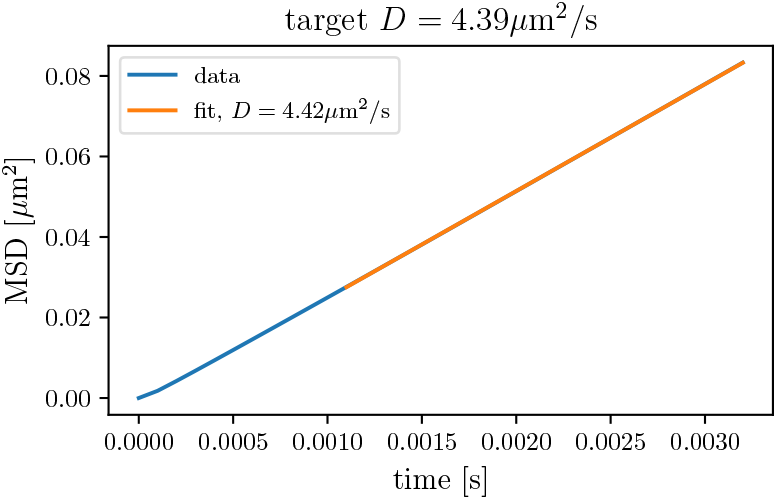
Phage mean squared displacement and diffusion coefficient fit.

### II. DETERMINATION OF INFECTION RATE

Figure S3 shows exemplary encounter data for a single simulation and the fit that is used to determine the infection rate.

**FIG. S3.**
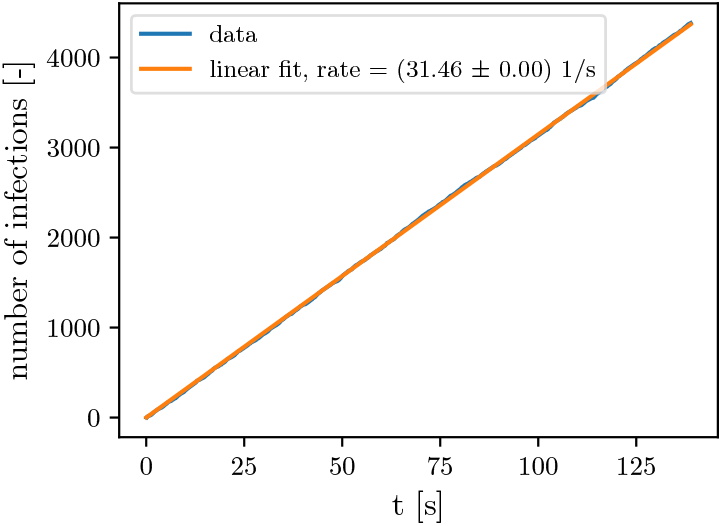
Number of infection vs time for one simulation with hydrodynamics, *k*^*att*^ *→ ∞* and *v*_*swim*_ = 100 μm s^*−*1^

### III. EXCLUSION OF FINITE SIZE AND DENSITY EFFECTS

In fig. S4 we show that the infection rate is independent of the box size and independent of the phage density.

**FIG. S4.**
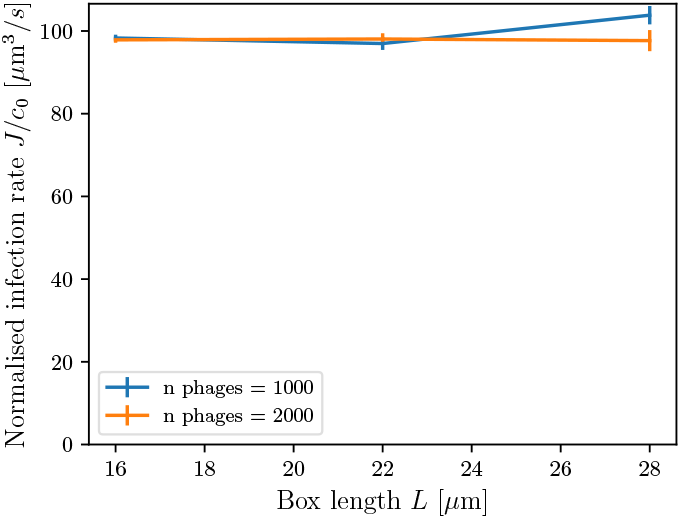
Infection rate as a function of simulation box length for different numbers of phages. Simulations performed with hydrodynamics at *v*_swim_ = 25 μm s^*−*1^ and *k*^att^ *→ ∞*.

### IV. DESCRIPTION OF SI MOVIE

`simulation_setup.mp4:` A pusher bacterium swimming at velocity *v*_swim_ = 25 μm s^*−*1^ in a periodic simulation box of dimension (25 μm)^3^ with 100 phages. The size of the phages is increased by a factor of 3 for better visibility.

### V. FINITE ATTACHMENT RATE

Figure S5 and fig. S6 show the results for infection rate and location on the body and flagellum, respectively. The data for *k*^att^ → 0, ∞ are the same as in the corresponding figures in the main text. For intermediate attachment rates, the curves show the same qualitative behaviour as in the two limiting cases.

### VI. PULLER SWIMMERS

Figure S7 and fig. S8 show results for infection rate and position for the body and flagellum, respectively. To obtain this data, puller type swimmers with the flagellum in front of the cell body were simulated.

Phages in the swimming direction are taken up by the flagellum before they can reach the front of the cell body. Therefore, in simulations without hydrodynamics, the main mechanism of infection rate increase on the cell body – uptake of more phages in the front – is strongly reduced. As a result, the infection rate increase is much smaller than for pushers and is also not captured by the cross sectional model.

When hydrodynamic interactions are considered, there is almost no increase in infection rate on the cell body regardless of *k*^att^. For the pusher, the infection rate increases with *v*_swim_ because the region with 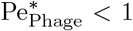 shrinks around the forward end of the cell body and phages have to cross less distance by diffusion. For the puller, the shape of 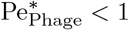 around the forward end of the cell body is roughly independent of the swim speed as it is mainly determined by the details of the gap between cell surface and the propelling part of the flagellum. Therefore, the only increase of infection rate comes from phages that arrive from the sides, leading to the small influence of motility. Unsurprisingly, the model of Berg et al. cannot be employed for puller bacteria and flagellotropic phages, because now the flagellum is the main collector of phages instead of the cell body.

The dependence of the infection location on the cell body is qualitatively the same as in the pusher case, except that the curves are shifted to negative values at *v*_swim_ = 0 because of the sink of phage concentration now in front of the cell body.

For the flagellum, the qualitative observations and explanations for the behaviour of infection rate and location are the same as in the pusher case.

**FIG. S5.**
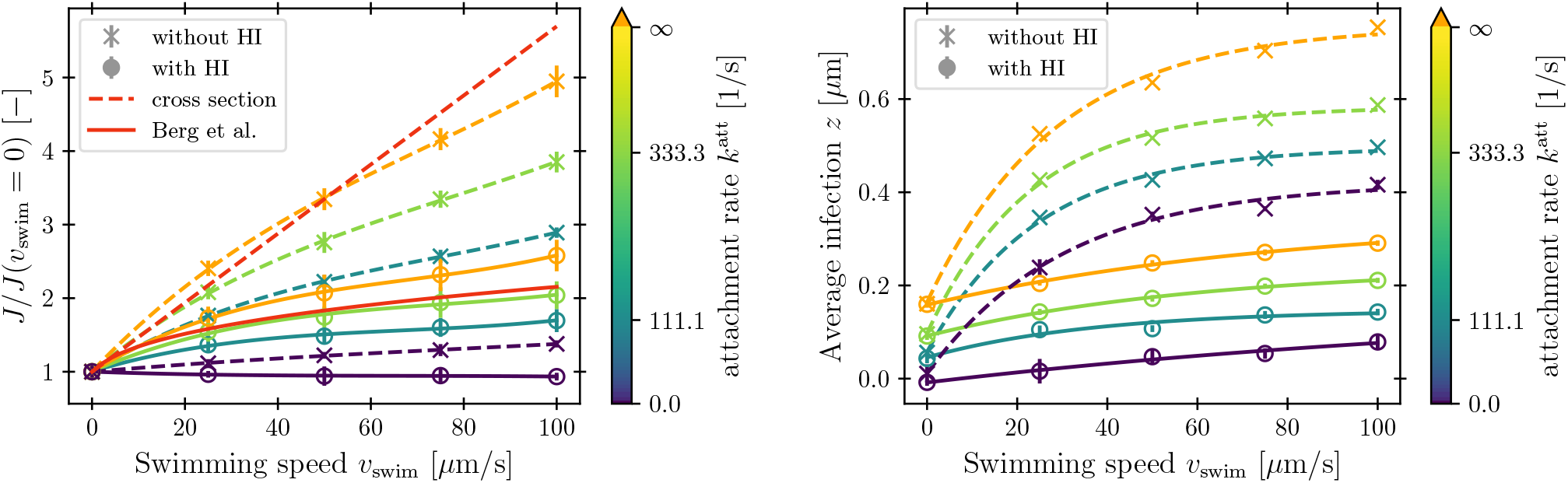
**Pusher** bacterium: Infection rate and positions on the **cell body** for finite attachment rate. Polynomial/exponential fits are shown to guide the eye.

**FIG. S6.**
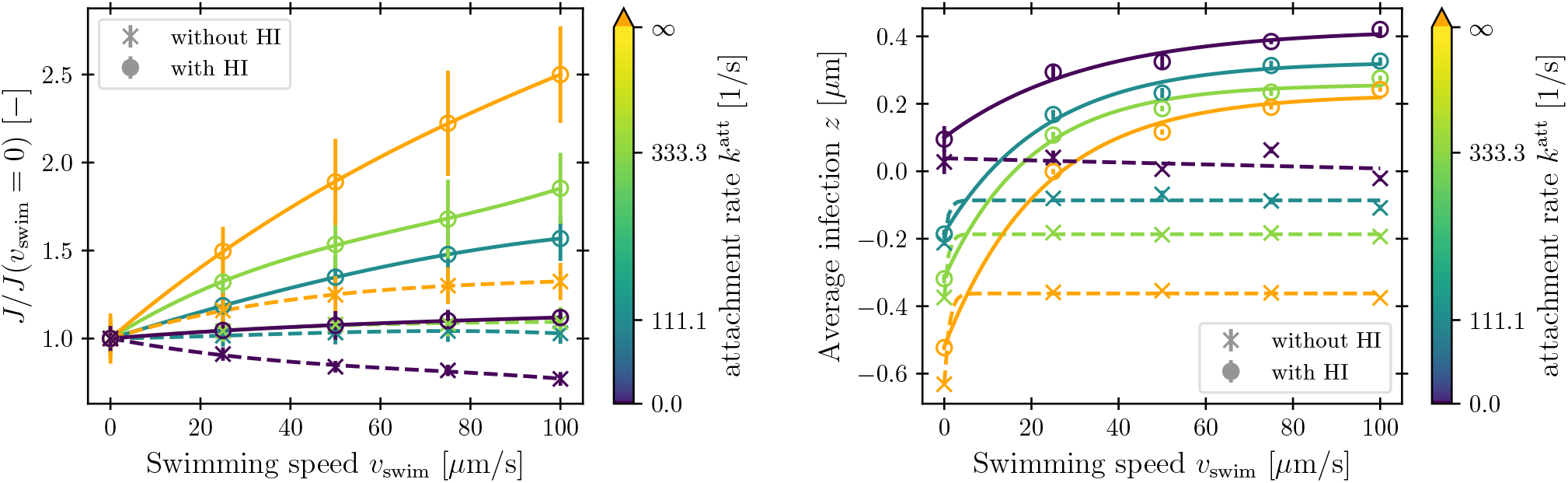
**Pusher** bacterium: Infection rate and positions on the **flagellum** for finite attachment rate. Polynomial/exponential fits are shown to guide the eye.

**FIG. S7.**
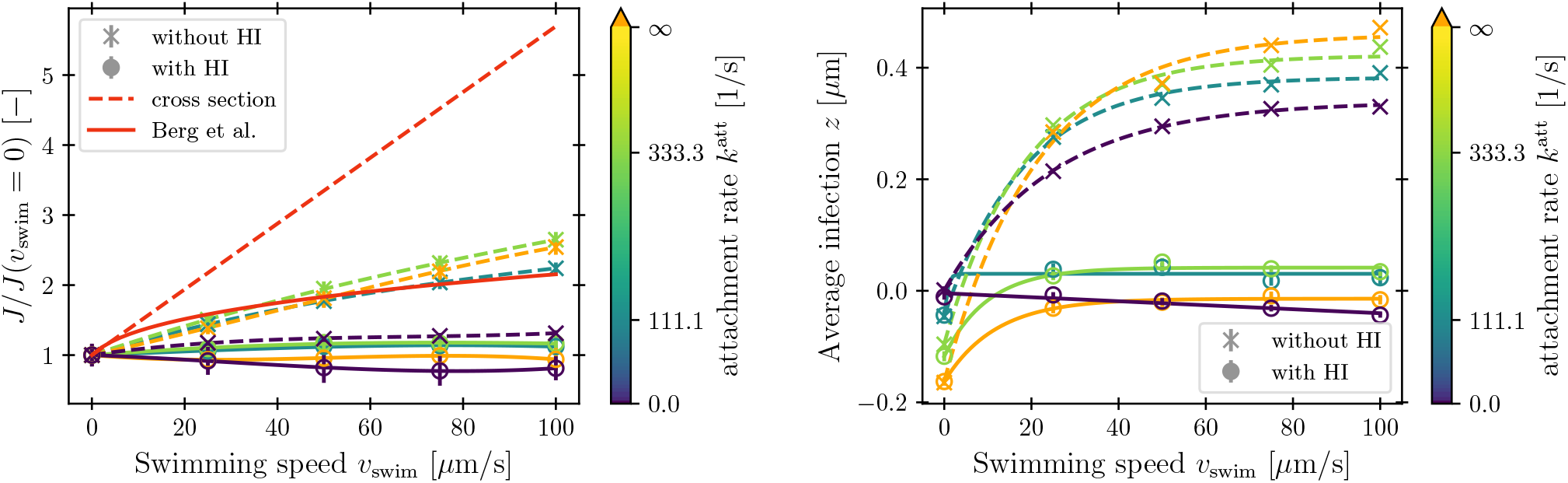
**Puller** bacterium: Infection rate and positions on the **cell body** for finite attachment rate. Polynomial/exponential fits are shown to guide the eye.

**FIG. S8.**
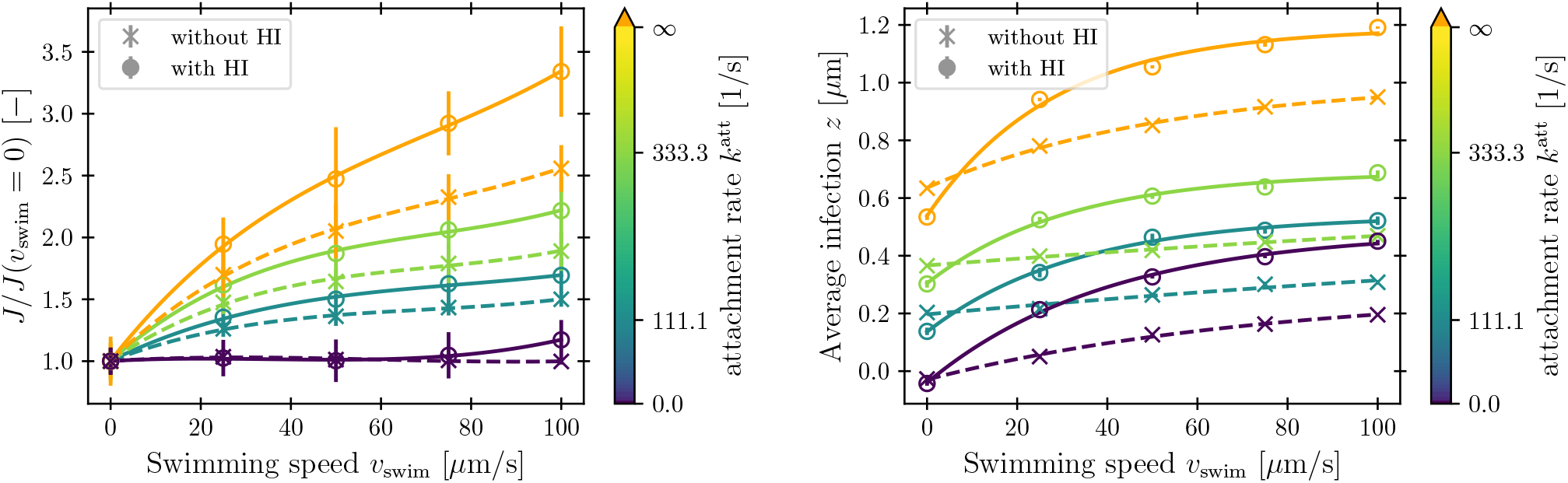
**Puller** bacterium: Infection rate and positions on the **flagellum** for finite attachment rate. Polynomial/exponential fits are shown to guide the eye.

